# Regenerative Neural Stem Cell Therapy Improves Multidomain Neurological Deficits after Traumatic Brain Injury in Nonhuman Primates

**DOI:** 10.64898/2026.08.20.745982

**Authors:** Marco Arredendo, Etienne W. Daadi, Elyas S. Daadi, Thomas Oh, Joshua Karam, Hooman Sadighian, Rebecca Nishi, Brian J. Cummings, Marcel M. Daadi

**Author notes:** These authors secured funding for the study (Brian J. Cummings, Marcel M. Daadi). **Correspondence:** M.M.D.

## Abstract

Traumatic brain injury (TBI) produces persistent multidomain disability spanning motor, cognitive, emotional and sleep–wake function, with no approved restorative therapy. Here, we tested pd.S6.133.hNSC, a cryopreserved, GMP-like human neural stem cell (hNSC) product derived from Shef-6 and FACS-sorted on CD133+/CD34−, in a randomized dose-ranging study in common marmosets subjected to controlled cortical impact (n = 18). Seven weeks after injury, animals received MRI-guided stereotactic transplantation into perilesional cortex bilaterally under tacrolimus immunosuppression, with either vehicle or pd.S6.133.hNSC at 1×10^6^ or 5×10^6^ cell dose. At 3 months post-transplantation, 5x10^6^ dosage improved executive and problem-solving performances (Object Retrieval Task with Barrier Detour), gait dynamics (CatWalk assay), anxiety-like behavior (Human Intruder Test), and actigraphy-derived sleep–wake and circadian rhythm measures relative to vehicle and 1x10^6^ dose. Longitudinal 7T MRI demonstrated a dose-dependent reduction in lesion volume and preservation of corpus callosum white matter volume in the 5x10^6^ group. Transplantation was well tolerated, with no observed adverse events across 1,197 cumulative post-transplant animal-days. Histopathology at 3 months post-transplantation in NHPs showed engraftment without tumor formation or abnormal tissue overgrowth. These findings support the safety and multidomain efficacy of a cryopreserved hNSC product in a nonhuman primate TBI model and inform translational development toward first-in-human testing with clinically aligned endpoints.

## Introduction

Traumatic brain injury (TBI) is a major global driver of death and long-term disability, with survivors frequently experiencing persistent multidomain impairments that extend far beyond focal neurological deficits. In addition to motor dysfunction, individuals commonly develop enduring executive and memory problems, affective disturbance, anxiety and depression, disrupted sleep–wake architecture and fatigue, and diminished community participation and quality of life. These symptoms can persist for years, often with incomplete recovery even after intensive rehabilitation, and they contribute substantially to socioeconomic burden and caregiver strain [1–32]. Recent consensus-driven efforts to develop standardized outcome sets for moderate-to-severe TBI have further highlighted the limitations of relying predominantly on global neurological scales and the importance of incorporating patient-centered outcomes across multiple functional domains into therapeutic trials [33]. Recent systematic reviews and meta-analyses similarly underscore the high prevalence and persistence of sleep disturbance, cognitive impairment and neuropsychiatric sequelae following TBI, reinforcing the need to evaluate therapeutic benefit beyond a single neurological endpoint [34–36].

Despite extensive preclinical and clinical development efforts—including more than a hundred interventional trials—no therapy has consistently improved the complex constellation of chronic deficits after moderate to severe TBI, nor the evolving pathology that includes axonal injury, white matter degeneration, maladaptive synaptic remodeling and chronic neuroinflammation [37–41]. This translational gap reflects, in part, the heterogeneity of injury patterns and the multifocal biology of secondary injury cascades that may continue for months to years after the initial mechanical insult [25–28, 42].

The most widely tested pharmacological approaches have focused on acute neuroprotection yet have largely failed to produce durable improvements in function across clinical domains. Even interventions with physiological plausibility, such as therapeutic hypothermia, have not been adopted as standard of care for most patients owing to variable trial outcomes, logistical constraints and uncertain benefit–risk profiles across patient subgroups [37]. Meanwhile, clinical and epidemiological work has sharpened the understanding that chronic TBI is not a static lesion but an evolving disorder with progressive network dysfunction. Diffuse axonal injury and demyelination disrupt long-range connectivity, particularly affecting frontal–striatal and interhemispheric pathways that support executive function, gait automaticity, emotional regulation and sleep stability [24–26, 43–45]. Recent network-level meta-analytic evidence further demonstrates widespread abnormalities of functional connectivity across TBI severity, supporting a framework in which clinical deficits emerge from disruption of interconnected large-scale systems [45]. In parallel, chronic neuroimmune activation and glial scarring can limit endogenous repair and perpetuate synaptic and white matter abnormalities [25, 26, 46]. These convergent mechanisms motivate a reparative strategy capable of engaging multiple pathological nodes rather than a single molecular target.

Cell-based therapies offer a potential route to multi-mechanistic repair. However, much of the clinical experience in TBI has come from non-neural products (for example, autologous bone marrow mononuclear cells or mesenchymal stromal/stem cells (MSCs)), delivered intravenously or by intracranial implantation. These studies have generally established feasibility and short-term safety but have shown variable or modest efficacy and often emphasize limited endpoint domains (e.g., motor outcomes) without robust evaluation of cognition, affect and sleep domains that patients prioritize and that frequently drive long-term disability [38–41, 47]. For example, the STEMTRA program of modified MSC implantation in chronic TBI provided evidence that stereotactic delivery is feasible and may improve motor measures but did not directly address circuit-level replacement or broader neurobehavioral recovery [47]. Together, these trials underscore both the promise and limitations of current cellular approaches and highlight the need for products with neural lineage relevance, rigorous dose-ranging, clinically aligned delivery, and multidomain outcome measures.

Human neural stem cells (hNSCs) offer a mechanistically compelling alternative because they can (i) generate neurons and glia relevant to cortical circuitry, (ii) support remyelination through oligodendrocyte lineage potential, and (iii) provide trophic and immunomodulatory signals that may attenuate chronic inflammation and promote endogenous plasticity [17, 42, 48–50]. Yet, translation of hNSC therapeutics has been constrained by challenges extending beyond biological efficacy, including variability in starting and ancillary materials, manufacturing changes, lot-to-lot consistency, product identity, potency, formulation and cryopreservation [51–53]. Recent regulatory guidance has placed particular emphasis on demonstrating manufacturing control, comparability and potency assurance for cellular and gene therapy products, underscoring that reproducible clinical translation requires a well-defined product rather than efficacy demonstrated with a nominally similar cell population [51–53].

To address these translational hurdles, we developed pd.S6.133.hNSC, a process-development (pd) “GMP-like” hNSC product derived from the Shef6 human embryonic stem cell line and enriched for CD133^+^/CD34^−^ expression via FACS, and manufactured as a cryopreserved, release-tested clinical candidate. In rodent TBI models, a research-grade, xeno-free version of this lineage has demonstrated consistent engraftment and tri-lineage differentiation, reduced neuroinflammation, improved cognitive performance and evidence of synaptic integration, supporting biological plausibility and cross-study reproducibility [42, 48–50]. The transition from this research-grade product to a defined cryopreserved clinical candidate was designed to address a central translational challenge in cell therapy: preservation of product identity and biological activity across manufacturing and formulation changes [51–53]. However, primate validation is essential before first-in-human testing because primate neuroanatomy, white matter organization, immune response and behavioral repertoire more closely approximate human disease.

Here we report an IND-enabling, dose-ranging study of pd.S6.133.hNSC in a nonhuman primate (NHP) controlled cortical impact model using the common marmoset. We transplanted cryopreserved pd.S6.133.hNSC at two doses (1 million and 5 million cells) into MRI-defined cortical targets 7 weeks after injury under clinically aligned tacrolimus immunosuppression. We then assessed recovery across endpoints chosen for direct clinical mapping: executive planning and fine visuomotor control (Object Retrieval Task with Barrier Detour), gait dynamics (CatWalk), anxiety-like behavior (Human Intruder Test), and sleep–wake rhythm and activity fragmentation (Actiwatch), alongside longitudinal MRI measurement of lesion evolution and white matter volume. We show dose-dependent improvements across motor and non-motor domains and structural repair signatures, supporting the premise that a defined, cryopreserved hNSC product can safely engage multiple symptom dimensions in a primate model of TBI.

## Results

### Human neural stem cells pd.S6.133.hNSC exhibit neural lineage potential and meet manufacturing release criteria

The pd.S6.133.hNSCs are CD133⁺/CD34⁻ sorted neural precursors derived from the Shef6 human embryonic stem cell line [54]. In this study the hNSCs were manufactured under process-development (pd) current good manufacturing practices (cGMP)-like conditions and cryopreserved at passage 7. The pd.S6.133.hNSCs expressed the neural stem cell marker nestin and retained the capacity to differentiate into βTubIII⁺ neurons, GFAP⁺ astrocytes, and Olig2⁺ oligodendrocytes, confirming multipotent neural lineage potential (Fig. 1).

**Figure 1:**
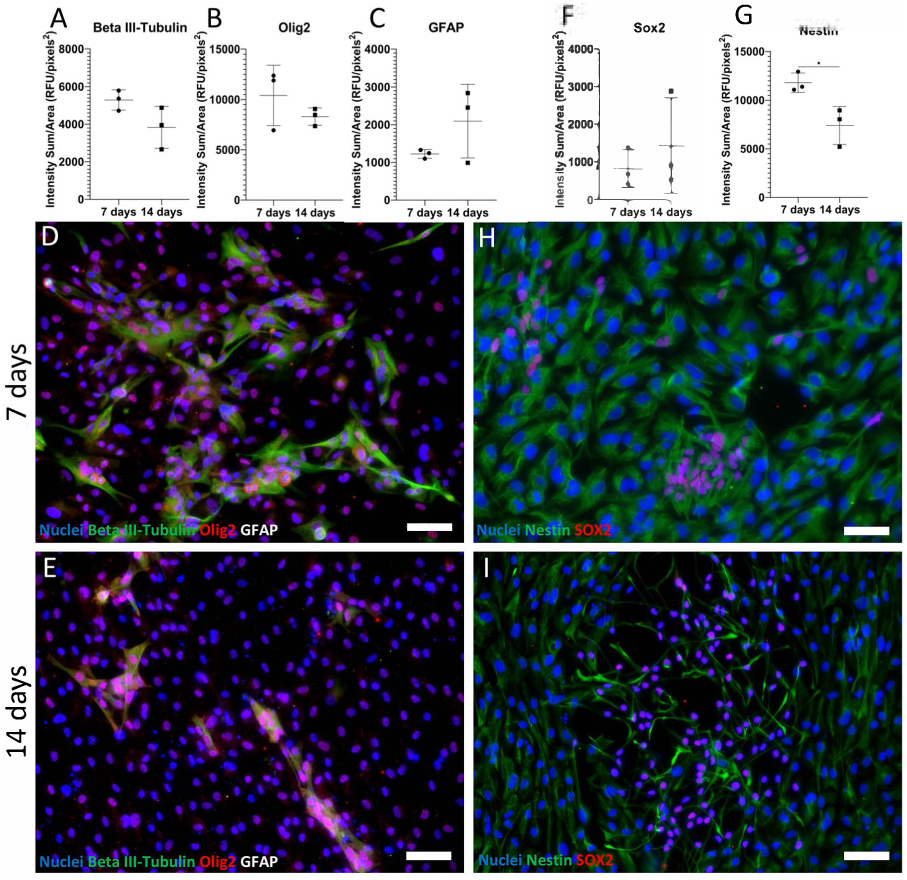
Pd.S6.133.hNSC differentiate into the three neural lineages, neurons, astrocytes and oligodendrocytes. **Pd.S6.133.hNSC** were thawed in growth media and seeded 10000 cells/well in an 8-well chamber slide. After 24 hours, the media was changed to differentiation media. After 7 and 14 days, cells were fixed and immunostained to assess expression of the differentiation markers Beta III-tubulin, Olig2, and GFAP (A-E) and the neural stem cell markers Sox2 and Nestin (F-I). Hoescht was used to counterstain nuclei (blue). The quantitative analysis showed as expected a significant decrease in the neural precursor marker nestin expression from 7 to 14 days of differentiation. Scale bars are 20 µm in D, E, H and 50 µm in I.

The production lot used for the present study pd.S6.133.hNSCs met all release criteria, including ≥85% CD133⁺/CD34⁻ purity and ≥70% post-thaw viability, demonstrating robustness of the manufacturing and cryopreservation process. The cell product exhibited normal karyotype and passed genome integrity, sterility, mycoplasma, and endotoxin testing, supporting suitability for translational development.

### Pd.S6.133.hNSC, transplantation is well tolerated in a nonhuman primate model of traumatic brain injury

To evaluate the safety and therapeutic potential of pd.S6.133.hNSCs, we transplanted cryopreserved hNSCs into a NHP model of traumatic brain injury using the common marmoset. Marmosets subjected to controlled cortical impact injury (CCI) [55, 56] (n = 18) were randomly assigned to receive vehicle, a low dose (1 × 10⁶ cells), or a high dose (5 × 10⁶ cells) of pd.S6.133.hNSCs seven weeks after injury.

Cryopreserved pd.S6.133.hNSCs were thawed and formulated in Isolyte at 100,000 cells/µl then stereotactically delivered into three cortical targets within the lesion site in the primary motor cortex, defined using MRI-based lesion mapping [57]. Injection sites were spaced along the rostrocaudal axis to maximize coverage of the injury cavity. Tacrolimus immunosuppression was initiated 2 days before transplantation and maintained until necropsy.

Animals were monitored longitudinally for adverse events including changes in weight, appetite, wound healing, and behavioral indicators of distress. No procedure- or treatment-related adverse events were observed following transplantation. Body weight remained stable throughout the study period, with no significant differences between groups over the three-month survival period (Data not shown). Across all animals, the study accumulated 1,197 marmoset-days of survival following intracranial pd.S6.133.hNSC transplantation under immunosuppression without evidence of treatment-related complications.

These findings support the tolerability of both low- and high-dose pd.S6.133.hNSC transplantation in the injured primate brain.

### Transplantation of pd.S6.133.hNSC improves executive function and sensorimotor planning after TBI

Executive function and visuomotor coordination were assessed using the Object Retrieval Task with Barrier Detour (ORTBD), a behavioral paradigm sensitive to dysfunction of frontal and motor cortical circuits. In this task, animals must plan and execute a detour movement to retrieve a reward from a transparent box with an obstructed opening [58, 59]. marmosets treated with the high-dose group (5 × 10⁶ cells) demonstrated significantly improved task performance compared with vehicle-treated animals. Specifically, high-dose treated animals required fewer reach attempts to retrieve the reward (Fig. 2A) and exhibited significantly fewer barrier-reaching errors (Fig. 2B).

**Figure 2:**
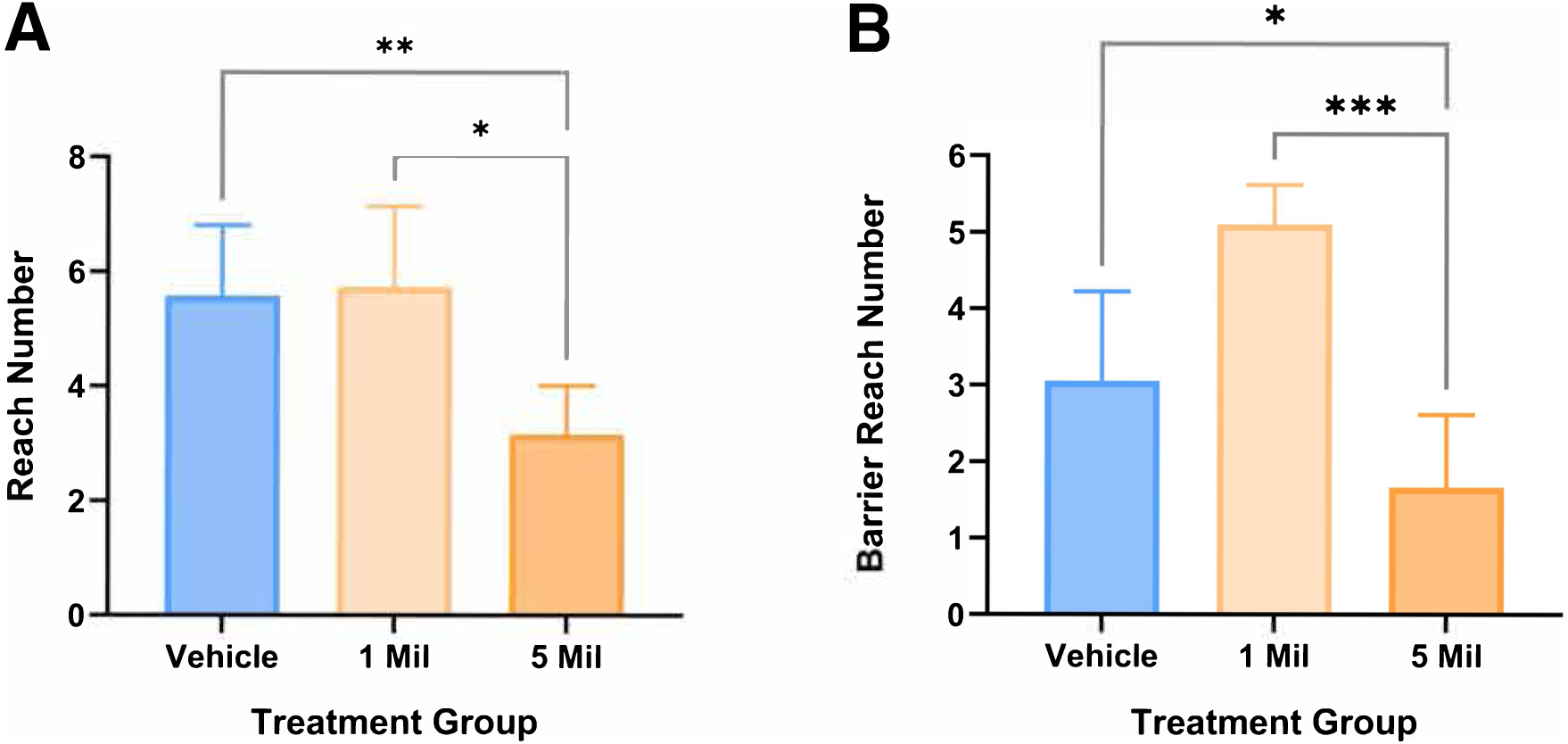
Higher dose pd.S6.133.hNSC improves executive function and problem-solving after TBI. A) Total reach number in ORTBD at 3 months post-transplantation. B) Barrier reach number. Groups include vehicle, 1 Mil (1×10^6^) cell dose and 5 Mil (5×10^6^) cell dose. Data are mean ± s.e.m. One-way ANOVA with Newman–Keuls multiple comparisons; *P < 0.05.

These findings indicate that high-dose pd.S6.133.hNSC transplantation improves executive planning and visuomotor coordination following TBI.

### Pd.S6.133.hNSC transplantation restores circadian activity and sleep architecture

Sleep disturbance and altered circadian activity are common sequelae of TBI. To assess whether pd.S6.133.hNSC treatment influences these parameters, general animal activity and sleep patterns were continuously monitored using collar-mounted actigraphy devices, the Actiwatch Mini [59].

Animals receiving the high-dose treatment exhibited increase in diurnal activity (Fig. 3A). In parallel, high-dose treated animals showed reduced nocturnal activity compared with both vehicle and low-dose groups (Fig. 3B) and longer nocturnal immobility periods, consistent with improved sleep consolidation (Fig. 3C).

**Figure 3:**
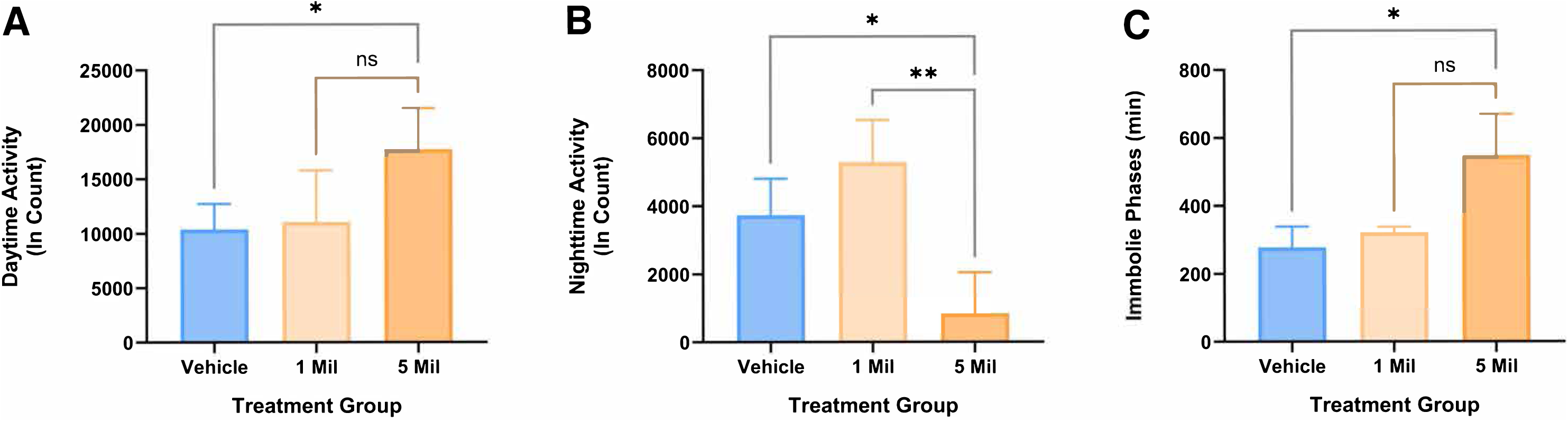
Pd.S6.133.hNSC improves rest–activity and sleep quality after TBI. Actigraphy outcomes at 3 months post-transplantation: A) diurnal activity, B) nocturnal activity and C) nocturnal immobility. Groups include vehicle, 1 Mil (1×10^6^) cell dose and 5 Mil (5×10^6^) cell dose. Data are mean ± s.e.m. One-way ANOVA with Newman–Keuls multiple comparisons; *P < 0.05.

These results suggest that pd.S6.133.hNSC transplantation promotes normalization of sleep–wake rhythms and daytime activity levels following TBI.

### Pd.S6.133.hNSC transplantation reduces anxiety-like behavior in TBI marmosets

Neuropsychiatric symptoms such as anxiety are frequent long-term consequences of TBI. Anxiety-like behavior was evaluated using the Human Intruder Test, which measures defensive behavioral responses to a novel human presence [60].

Marmosets treated with pd.S6.133.hNSC exhibited reduced anxiety-like responses compared with vehicle-treated animals. In particular, animals receiving the high-dose treatment (5 × 10⁶ cells) spent significantly less time in elevated defensive positions within the cage during intruder exposure (Fig. 4A). Additionally, the duration of head-bobbing behavior, an established indicator of anxiety in marmosets, was significantly reduced in the high-dose treatment group (Fig. 4B).

**Figure 4:**
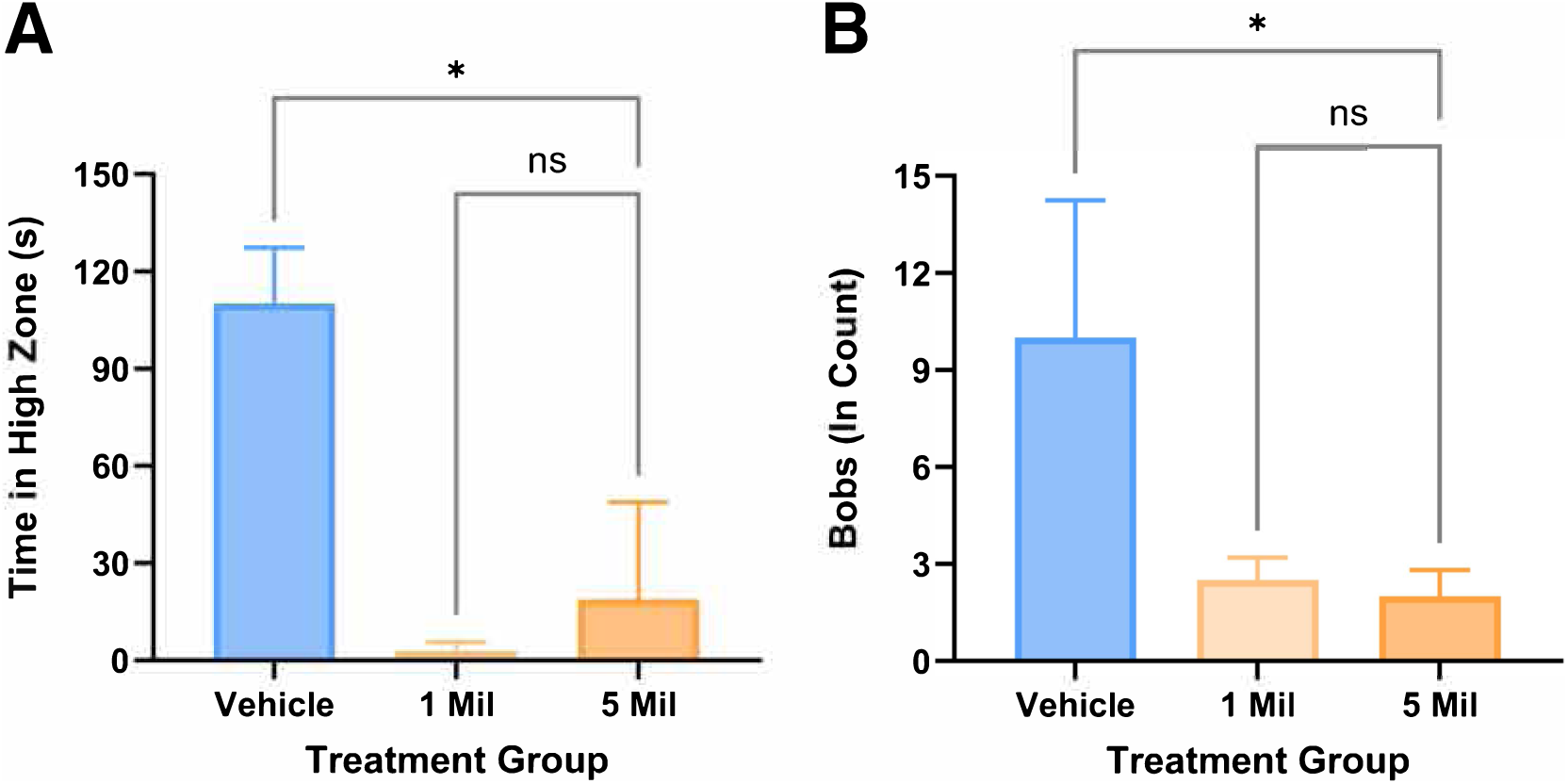
pd.S6.133.hNSC reduces anxiety-like behavior in the Human Intruder Test. A) Time spent in the high zone of the cage during intruder exposure. B) Head-bobbing duration. Data are mean ± s.e.m. One-way ANOVA with Newman–Keuls multiple comparisons; ***P < 0.001; *P < 0.05.

These results indicate that pd.S6.133.hNSC transplantation alleviates anxiety-like behaviors associated with TBI.

### Transplantation of pd.S6.133.hNSC improves gait and locomotor coordination

Motor coordination and locomotion were assessed using the CatWalk XT automated gait analysis system. This platform quantifies paw placement and gait dynamics during voluntary locomotion. Each animal completed 9 runs (3 trials × 3 runs) on the walkway. animals receiving the high-dose pd.S6.133.hNSC treatment demonstrated a restoration of the baseline of footfall patterns during locomotion (Fig. 5A) and exhibited a significant increase in paw swing duration compared with vehicle-treated animals (Fig. 5B). Increased paw swing reflects improved dynamic limb movement during gait cycles, suggesting enhanced locomotor coordination.

**Figure 5:**
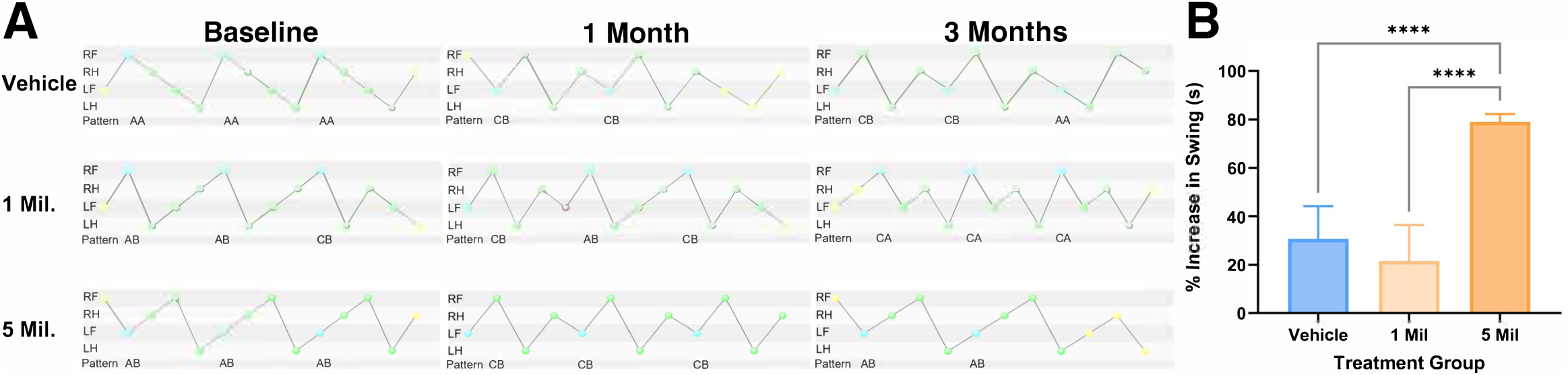
Higher-dose pd.S6.133.hNSC restores footfall patterns and improves gait dynamics. **A)** CatWalk XT analysis showing patterns of footfall sequence during locomotion. Colored spheres indicate phases of the pattern, blue = start, green = part of the pattern, red = not part of the pattern and yellow = not taken into account. The high dose group (5 Mil) shows restoration of the normal gait, with sequential use of paws pattern AA or AB: AA Pattern: RF > RH > LF > LH; AB Pattern: LF > RH > RF > LH). CB Patterns: abnormal, ataxic gait, with diagonal use of paws. (LF > RF > LH > RH). CA Pattern: abnormal gait (e.g. RF->LF->RH->LH). **B**) CatWalk XT analysis showing percentage paw swing at 3 months post-transplantation. Each animal performed 9 runs (3 trials × 3 runs). Data are mean ± s.e.m. One-way ANOVA with Newman–Keuls multiple comparisons; ***P < 0.001.

These findings indicate that pd.S6.133.hNSC treatment improves sensorimotor integration and gait function following traumatic brain injury

### Pd.S6.133.hNSC transplantation reduces lesion volume and restores white matter integrity

To determine whether functional improvements were associated with structural brain repair, lesion volume and white matter integrity were assessed longitudinally using MRI. Analysis of serial T2-weighted MRI scans revealed that animals receiving the high-dose pd.S6.133.hNSC treatment exhibited significantly reduced lesion volumes three months after transplantation compared with vehicle-treated animals (Fig. 6B). In addition, pd.S6.133.hNSC-treated animals demonstrated increased white matter volume within the corpus callosum relative to vehicle-treated controls (Fig. 6C).

**Figure 6:**
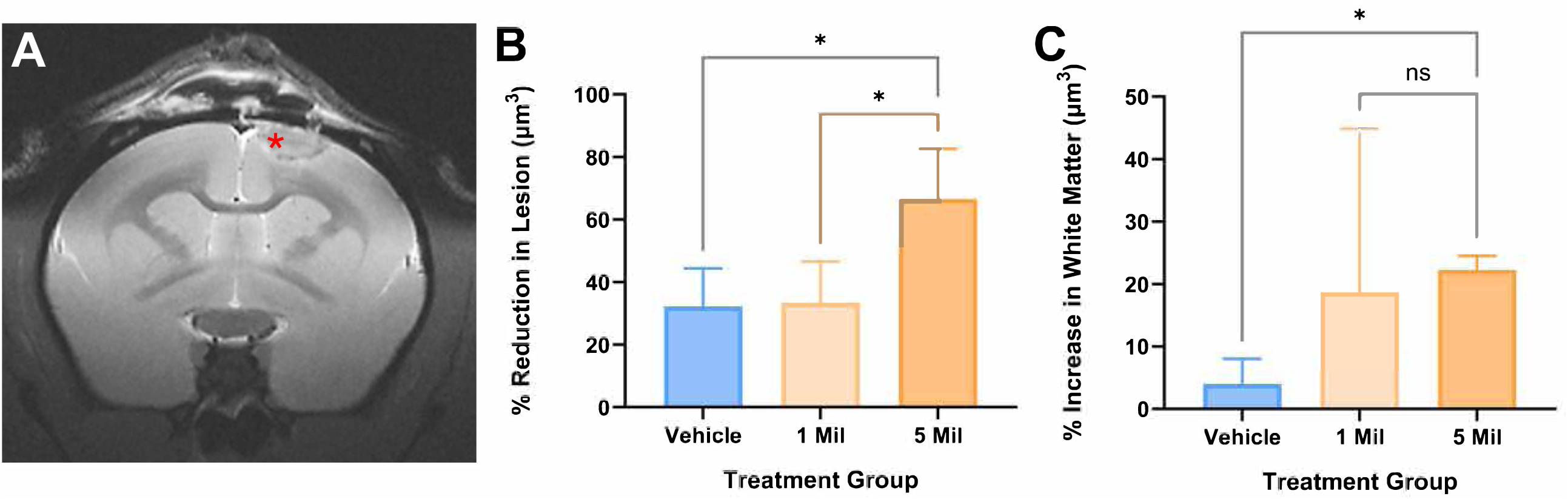
7T MRI reveals dose-dependent reduction in lesion volume and increased corpus callosum white matter volume. **A)** T2-weighted coronal MR scan image showing the TBI (red star) in the cerebral cortex areas. **B**) Percent lesion volume reduction from pre-transplant to 3 months post-transplantation. **C**) Change in corpus callosum white matter volume over the same interval. Data are mean ± s.e.m. One-way ANOVA with Newman–Keuls multiple comparisons; *P < 0.05.

These imaging findings suggest that pd.S6.133.hNSC transplantation promotes structural repair of injured brain tissue, including preservation or restoration of white matter tracts.

### Pd.S6.133.hNSC grafts engraft without tumor formation and integrate within host tissue

Postmortem histological analysis was performed to evaluate graft survival, differentiation, and potential adverse tissue effects.

Hematoxylin and eosin (H&E) staining (Fig. 7A) revealed no evidence of tumor formation, teratoma development, or abnormal tissue structures in pd.S6.133.hNSC - treated brains. Histopathological findings such as hemosiderin deposition, macrophage infiltration, and astrocytosis were consistent with the original traumatic injury and were observed across experimental groups.

**Figure 7:**
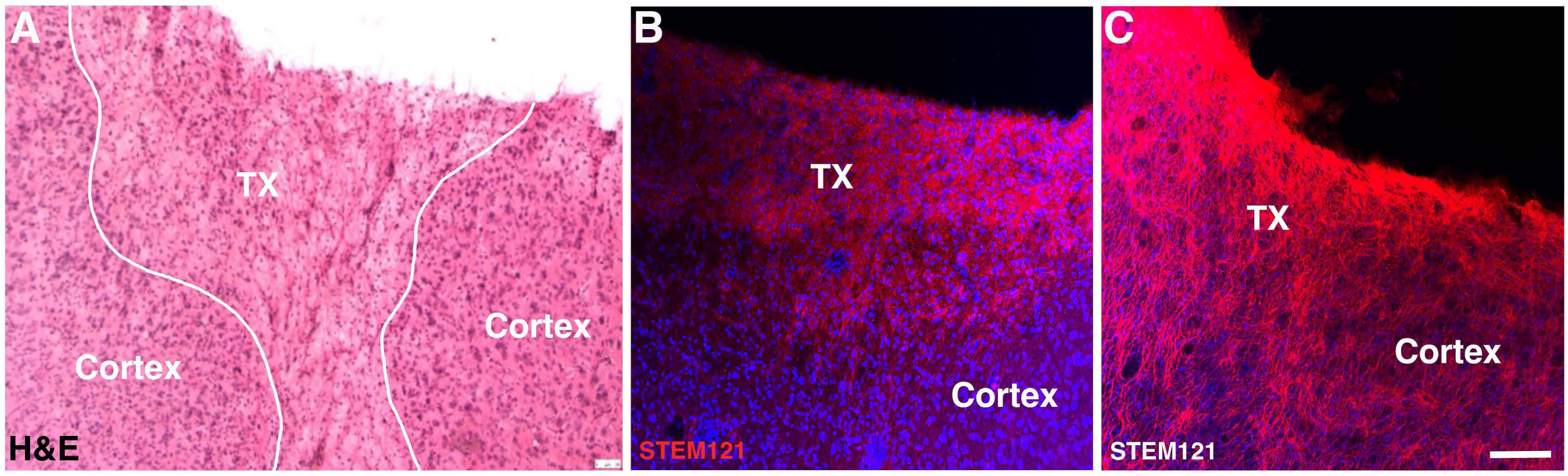
Pd.S6.133.hNSC engraft into the peri-lesional cortex. Representative coronal sections showing A) H&E staining of the grafted hNSCs in the cerebral cortex, with no abnormal structural changes. B) and C) the grafted hNSCs, immuno-stained with the human cell marker STEM121, seamlessly integrated in the TBI lesioned cortex of the marmosets at 3 months post-transplantation, showing migration of the hNSCs and process elaboration into the host peri-lesional cerebral cortex. Scale bars are 50 µm in A, B and 80 µm in C.

Immunohistochemical staining using the human-specific cytoplasmic marker STEM121 (Fig. 7B, C) confirmed the engraftment of pd.S6.133.hNSC within the injured cerebral cortex. Grafted cells exhibited structural integration within the cytoarchitecture of the cerebral cortex and were distributed throughout the lesion site without evidence of aberrant structural changes.

Together, these findings demonstrate that pd.S6.133.hNSCs survive and integrate within the injured NHP cerebral cortex without evidence of tumorigenicity.

## Discussion

We report for the first time, that transplantation of the GMP-compatible pd.S6.133.hNSC cellular product into a TBI marmoset model produces dose-dependent improvements across clinically salient domains including motor, executive, affective and sleep-related. These outcomes were accompanied by MRI signatures consistent with reduced lesion burden and preservation or augmentation of white matter volume. The convergence of behavioral and imaging outcomes strengthens the translational argument for pd.S6.133.hNSC and informs critical parameters for first-in-human development: dosing, target selection, safety monitoring and endpoint strategy.

### Multidomain functional recovery is a primary translational hurdle in chronic TBI

Human TBI is characterized by heterogeneous, often persistent impairments that extend beyond motor disability. Sleep fragmentation and circadian disruption after TBI are strongly linked to fatigue, mood symptoms, cognitive complaints and reduced rehabilitation gains [22, 27–32]. Anxiety and related stress phenotypes are common, worsen participation in therapy and social reintegration, and are associated with increased suicidality in some populations [1–3, 12, 61–66]. These clinical realities have not been consistently mirrored in preclinical development pipelines, which frequently prioritize short-term sensorimotor endpoints. A central strength of the current work is the explicit focus on domains that are highly prevalent and clinically disabling—executive planning, sleep–wake activity fragmentation, and anxiety-like behavior—alongside gait. In this model, high-dose of pd.S6.133.hNSC improved performance on ORTBD, normalized activity patterns suggestive of improved sleep quality, reduced anxiety-like responses in the Human Intruder Test, and improved gait dynamics. This multidomain efficacy profile aligns with the hypothesis that stem cell–based interventions may act through distributed network repair and modulation rather than a single localized mechanism.

### Positioning relative to existing TBI cell therapy trials

Clinical trials using autologous bone marrow mononuclear cells and MSC-derived approaches have provided important safety precedent and helped refine delivery and trial conduct in TBI [38–41, 47]. However, these products are not intrinsically neural and are generally thought to act via transient immunomodulation and trophic signaling; their capacity to meaningfully support remyelination, synaptic integration or durable circuit reconstruction remains uncertain. The post hoc analyses from stereotactic MSC implantation trials suggest possible motor improvements [47], but broader cognitive, affective and sleep endpoints have been limited or absent, despite their centrality to patient burden. Our data support a rationale for deploying a neural lineage-specified product that may better match the biology of cortical and white matter injury, particularly in chronic stages characterized by disconnection and demyelination [25, 26].

Importantly, the present study also emphasizes dose dependence: the 5 million cell dose consistently outperformed the 1 million-dose across outcomes, suggesting that insufficient dosing may be one contributor to variability in earlier translational efforts.

### Structural repair signatures: lesion evolution and white matter volume

Longitudinal MRI showed reduced lesion volume and increased corpus callosum white matter volume in the high-dose pd.S6.133.hNSC group. Although volumetric measures do not identify the cellular mechanisms underlying these structural changes, they provide objective, clinically translatable biomarkers for longitudinally tracking injury evolution and progressive neurodegeneration after TBI. Serial volumetric MRI studies in human TBI have documented progressive regional brain and white matter loss over months to years, including involvement of the corpus callosum, and MRI-derived volumetric measures have been associated with injury severity and long-term functional outcome [67–70]. Quantitative MRI has also been incorporated as a structural endpoint in interventional TBI studies, including a randomized cell-therapy trial in which preservation of white matter volume and corpus callosum microstructure were evaluated longitudinally [71].

White matter preservation is particularly relevant to TBI because traumatic axonal injury and subsequent alterations in white matter integrity are major contributors to persistent cognitive dysfunction, with diffusion-based measures of white matter injury associated with slowed information processing and impaired executive function [72–74]. The corpus callosum is especially vulnerable to traumatic axonal injury and undergoes progressive macrostructural and microstructural changes after moderate-to-severe TBI [68, 69, 75]. Accordingly, the increased corpus callosum white matter volume observed following pd.S6.133.hNSC treatment is consistent with structural preservation or repair but does not, on its own, establish remyelination, axonal regeneration or other specific cellular mechanisms. This finding is nevertheless consistent with prior rodent studies of Shef6-derived hNSCs, in which transplanted cells survived and differentiated into neural lineages, including oligodendrocyte-lineage cells, and were associated with functional recovery and modulation of the injury environment [48–50].

Future studies incorporating diffusion MRI, including DTI and more advanced models such as NODDI, together with myelin-sensitive imaging such as myelin water fraction, could help distinguish changes in bulk white matter volume from alterations in axonal integrity, neurite density and myelin content [76–79]. Diffusion and myelin-water imaging have demonstrated sensitivity to longitudinal white matter abnormalities after human TBI, including corpus callosum pathology, and several imaging measures have been associated with injury severity, cognitive performance and clinical outcome [73, 76–79]. Thus, although the present MRI findings should not be interpreted as direct evidence of a specific repair mechanism, they provide a translational bridge between the structural effects of pd.S6.133.hNSC observed in this NHP model and quantitative imaging biomarkers that can be incorporated into the clinical development and future evaluation of pd.S6.133.hNSC in patients with TBI

### Mechanistic considerations: beyond cell replacement

Pd.S6.133.hNSC likely exerts therapeutic effects through multiple, potentially synergistic mechanisms. Prior rodent work with the lineage demonstrated tri-lineage differentiation, synaptic integration of graft-derived neurons and attenuation of microglial activation and astroglial reactivity [42, 48–50]. In the primate brain, similar mechanisms could contribute to functional recovery: (i) replacement of lost neural elements in peri-lesional cortex, (ii) support of remyelination through oligodendroglial differentiation or trophic support of endogenous oligodendrocyte progenitors, (iii) paracrine modulation of neuroimmune signaling and vascular remodeling, and (iv) promotion of adaptive plasticity in spared networks.

The observed improvements across sleep, anxiety and executive performance are consistent with a network-level influence extending beyond the immediate lesion core [43, 80, 81]. However, the present behavioral and structural data do not permit definitive mechanistic attribution. Establishing how pd.S6.133.hNSC influences host brain repair will require quantitative histological analyses of graft fate and lineage composition, synaptic integration and proliferation, together with spatial transcriptomic and single-cell profiling of host immune and glial states, which have revealed marked regional and cell-type-specific responses after TBI [82–85]. Circuit-level electrophysiology and longitudinal functional imaging will also be important to determine whether cellular engraftment and structural preservation are accompanied by restoration or reorganization of neural activity and functional connectivity [80, 81, 85–87]. Such multimodal studies will be necessary to link graft-associated cellular and molecular changes to recovery across the cognitive, emotional and sleep–wake domains.

### Dose, timing, delivery and immunosuppression: practical implications for translation

This work provides several parameters relevant to IND planning. First, the lack of adverse events across 1,197 cumulative post-transplant survival days supports tolerability of stereotactic cortical transplantation under tacrolimus immunosuppression in this model, and the absence of tumor formation or gross tissue abnormalities is consistent with the product’s release criteria and prior teratoma risk assessments [54]. Second, treatment at 7 weeks post-injury targets a late-subacute to early-chronic transition period, after the immediate acute phase of injury but while secondary pathological processes, tissue remodeling and network reorganization remain active. The temporal boundaries between subacute and chronic TBI are not universally defined; however, longitudinal human and experimental studies demonstrate that neuroinflammation, structural degeneration, synaptic remodeling and neuroplastic responses continue for weeks, months and, in some cases, years after the initial injury [24, 46, 88, 89]. Thus, this treatment window may represent a clinically relevant opportunity to intervene after initial lesion formation while endogenous repair and remodeling processes remain ongoing, rather than assuming that the injured brain has entered a biologically static chronic state [24, 46, 88–90]. Third, MRI-guided targeting of peri-lesional cortex is consistent with a strategy of repairing distributed cortical–cortical connections and supporting white matter integrity rather than focusing exclusively on deep structures. These choices mirror practical constraints and opportunities in human stereotactic neurosurgery. Together, the timing of intervention, MRI-based targeting and stereotactic delivery paradigm provide a practical translational framework that can inform surgical planning, dose selection and longitudinal safety monitoring in the future clinical development of pd.S6.133.hNSC.

### Limitations

Several limitations should be addressed in subsequent studies. Sample size (Table 1), while appropriate for a primate dose-ranging proof-of-concept, limits subgroup analyses (e.g., sex as a biological variable). Behavioral assays in NHPs can be influenced by motivation and environmental factors, and future studies should include additional blinding safeguards. The study duration (3 months post-transplant) captures early functional recovery but not long-term durability, late adverse events, or chronic graft maturation. Mechanistic endpoints are currently incomplete and will be important for establishing the differentiation to various lineages with multiomics analysis and for correlating graft metrics with functional outcomes.

**Table 1:** Cross-species and cross-laboratory reproducibility of functional efficacy following S6.133.hNSC transplantation after traumatic brain injury. To evaluate the reproducibility of the S6.133.hNSC cell therapy platform and compare efficacy across studies, we computed Cohen’s D for behavioral effect sizes in our past three rodent studies [48–50] and for the present marmoset study. Across all four independent studies, transplantation of the S6.133.hNSC consistently produced ‘medium’ to ‘large’ effect sizes for behavioral measures of functional recovery. (GMP: good manufacturing practices, MWM: Morris Water Maze, NHP: nonhuman primate, NPR: novel place recognition).

| *Study* | *Species* | *S6.133.hNSC*<br>*Lot* | *Test* | *CohenD* | *Effect*<br>*Size* |
| --- | --- | --- | --- | --- | --- |
| *Ref. #: 50* | Rat | Research Grade | NPR Discrimination Index | 2.17 | Large |
| *Ref. #: 49* | Rat | Research Grade | NPR Discrimination Index | 0.66 | Medium |
| *Ref. #: 48* | Rat | Research Grade | MWM Acquisition | 0.87 | Medium |
| *Ref. #: 48* | Rat | Research Grade | MWM Reversal | 1.10 | Large |
| *Ref. #: 48* | Rat | Research Grade | MWM Combined Probes | 1.33 | Large |
| *Ref. #: 48* | Rat | Research Grade | Elevated Plus Maze | 1.32 | Large |
| *This study* | NHP | GMP-like | Object Retrieval Reach Number | 1.46 | Large |
| *This study* | NHP | GMP-like | Object Retrieval Barrier Reach | 2.82 | Large |
| *This study* | NHP | GMP-like | ActiWatch Nocturnal | 2.33 | Large |
| *This study* | NHP | GMP-like | ActiWatch Immobile | 2.22 | Large |
| *This study* | NHP | GMP-like | ActiWatch Diurnal | 3.14 | Large |
| *This study* | NHP | GMP-like | White Matter - MRI | 5.54 | Large |
| *This study* | NHP | GMP-like | Lesion Volume - MRI | 2.40 | Large |

### Outlook and clinical trial implications

The data support advancing pd.S6.133.hNSC toward first-in-human testing in chronic TBI with a clinical design that explicitly includes multidomain outcomes and biomarkers. Candidate endpoint strategy should combine motor assessments with validated cognitive/executive tasks, patient-reported sleep and mood outcomes, actigraphy, and MRI-based lesion and white matter measures. Given the apparent dose dependence, careful attention to dose scaling, injection number, and target coverage will be essential. The present results justify a Phase 1 trial prioritizing safety, feasibility, and biomarker engagement, while informing selection of secondary endpoints most likely to detect clinically meaningful benefit.

## Acknowledgements

This work was supported by The California Institute for Regenerative Medicine CIRM TRAN1-11548 and by NeoNeuron LLC.

## Author Contributions

BJC, MMD: conceptualized and designed the overall study.

MA, EWD, ESD, TO: conducted the behavioral testing, animal care, postmortem analysis

EWD, ESD, TO, MMD: performed the neurosurgical procedures for traumatic brain injury in nonhuman primates and the neural stem cell transplantation.

MA, EWD, ESD, TO, BJC, MMD performed data analysis.

HS: Performed cell preparations

MMD: directed and supervised the entire project, provided core infrastructure

MMD: wrote the first draft of the manuscript

MA, EWD, ESD, TO, JK, HS, RN, BJC, MMD provided comments and final approval of the manuscript.

## Materials and Methods

### Animals

Common marmosets (Callithrix jacchus; n = 18; 9 males, 9 females) were housed with environmental enrichment and provided a complete life-cycle commercial diet supplemented with fruits and vegetables and water ad libitum. Temperature and humidity were maintained within species-appropriate ranges. Animals were socially housed when compatible with study procedures carried out in strict accordance with the recommendations proposed in the Guide for the Care and Use of Laboratory Animals, National Research Council U. S. A. The protocols were approved by the Institutional Animal Care and Use Committee for The University of Texas Health San Antonio (UTHSA). All nonhuman primates held and used at UTHSA are maintained under conditions that meet USDA Animal Welfare Regulations, OLAW standards, and National Institute of Health (NIH) guidelines as stated in the *Guide for the Care and Use of Laboratory Animals* (81h Edition, 2010), NAS-ILAR recommendations, and AAALAC accreditation standards for these species. UTHSA is fully accredited by AAALAC International. The center promotes social housing caging, with structural complexities for environmental enrichment with the detailed observation of ongoing animal activities. The temperature inside animal quarters is maintained at 80°F and humidity (60%) suitable for marmosets. Animals are fed constant nutrition, complete life-cycle commercial monkey chows, supplemented daily with fruits and vegetables, and drinking water. All research activity has been conducted in accordance with the IACUC oversight process. We have resources and expertise to provide program administration, animal husbandry, clinical medicine, psychological well-being, facilities maintenance, animal records, and technical research support. All procedures were performed to minimize discomfort, distress, or pain. When necessary, sedation and anesthetic agents are used to render the animal unconscious and therefore insensate to handling, discomfort, or pain. Likewise, when necessary, analgesics are used to reduce any potential pain. All animals are enrolled in the environmental enrichment program. Enrichment provided to the animals consists of social contact, structural enrichment (e.g., perches, swings), manipulable enrichment (e.g., chew toys, balls), nutritional enrichment (e.g., fruit, grain), sensory enrichment (e.g., television, radio), and occupational enrichment (e.g., food puzzles). All enrichment provided is documented, and any deficiencies are addressed. We perform humane euthanasia of animals and in accordance with the professional principles and practices specified by the *American Veterinary Medical Association Guidelines for the Euthanasia of Animals: 2013 Edition.* Animals destined for euthanasia are injected intraperitoneally with sodium pentobarbital overdose (100 mg/Kg) followed by transcardiac perfusion with phosphate buffered saline and 4% paraformaldehyde for tissue processing.

### Controlled cortical impact injury

Animals were anesthetized with isoflurane and placed in a stereotaxic frame. A 6-mm craniectomy was performed over the right frontoparietal cortex to expose the dura. TBI was induced using an electronically controlled impactor (TBI-0310 device) with a 5-mm diameter tip at a velocity of 5.25 m/s and an impact depth of 2 mm. Food consumption, body weight, attitude, posture, excreta, appearance, and movements were monitored after all surgical procedures. If an animal experienced loss of appetite, an electrolyte replacement drink, soft foods and fruits were offered. Analgesics were given for 2 days. Antibiotics were administered for 5 days following the surgical procedure.

### Experimental design, randomization and blinding

At about 7 weeks post-TBI, animals were randomized into three groups (vehicle n=6 (M/F: 3/3), 1×10⁶ cell dose group n=4 (M/F: 2/2) and 5×10⁶ cell dose group n=8 (M/F: 4/4). Behavioral scoring and MRI volumetry were conducted by personnel blinded to group identity. Both sexes were included; the study was not powered to detect sex-by-treatment interactions.

### Pd.S6.133.hNSC hNSC product

Pd.S6.133.hNSC hNSCs are CD133+/CD34−enriched neural precursors derived from the Shef6 human embryonic stem cell line [54]. Cells were manufactured under GMP-like conditions and cryopreserved at passage P7. Lot release criteria we used included post-thaw viability ≥70%, CD133+/CD34− purity ≥85%, normal karyotype and negative sterility, mycoplasma and endotoxin tests.

### Immunosuppression and clinical monitoring

Tacrolimus was administered orally at 0.15–0.20 mg/kg per day starting 2 days before transplantation and maintained until necropsy. Animals were monitored for food intake, body weight, posture, appearance, excreta and movement.

### MRI acquisition and reconstruction

MRI was performed 1-month post-TBI and 3 months post-transplantation on a 7T Bruker Biospec system. DTI was acquired using a single-shot spin-echo EPI sequence. Structural imaging included multi-echo T2 scans with TE = 8.9, 26.69, 44.48 and 62.27 ms; TR = 3045.559 ms; RARE factor = 2; NEX = 8; refocusing angle = 180°; matrix = 128×128; field of view = 1.28×1.28 cm. Raw data were processed using MATLAB code to generate T2 output, and images were reconstructed to 256×256. Total scan time was approximately 10 minutes per subject. Lesion volume and corpus callosum white matter volume were quantified from serial coronal MRI images using ImageJ / Fiji as previously described [91, 92]. Regions were traced per slice and volumes calculated by summing areas multiplied by slice thickness. Quantification was performed blinded to treatment group.

### Stereotactic cell transplantation

Cryopreserved pd.S6.133.hNSC hNSCs were thawed and formulated in Isolyte at 100,000 cells/µl. MRI-derived lesion anatomy was used to define 3 cortical perilesional targets bilaterally (ipsilateral and contralateral to lesion) spanning the rostral, caudal and lateral boundaries of the TBI lesion. For each of the 6 trajectories per animal (3 ipsilateral and 3 contralateral to the lesion), volume per trajectory was scaled to deliver an equal share of the assigned dose at 100,000 cells/µl. Vehicle (Isolyte) was infused at the matched volume. Infusion rate was 1 µl/min via a controlled perfusion pump. After each injection, the cannula remained in place for 5 min and was then withdrawn at 1 mm/min to minimize reflux [57].

### Object Retrieval Task with Barrier Detour (ORTBD)

ORTBD was conducted as previously described [58, 59] with modifications for marmoset home-cage testing. Animals were acclimatized to the apparatus before testing. In each trial, animals retrieved a marshmallow reward from the open side of a transparent box; the opening orientation was randomized (left, right or toward the cage opening). Testing was performed for 3 consecutive days with 20 trials per day. Sessions were video-recorded and scored offline by blinded raters. Primary outcomes reported were total reach number and barrier reach number.

### Activity, sleep, and circadian rhythm analysis

Diurnal activity and sleep-wake patterns were monitored using Actiwatch Mini (Camntech) mounted on a collar [59]. Animals were acclimatized to the collar via progressive exposure sessions (15 min to 12 h). Actigraphy was recorded for 24 h on three separate days. Devices were placed at 08:30 in the morning and programmed to record from 09:00 for 24 h; data were downloaded via Sleep Analysis 7 software. Sleep analysis quantified sustained quiescence from 19:00 to 06:30 (∼11.5 h), corrected for individual sleep-onset timing to keep the analyzed sleep window consistent across animals. Nonparametric circadian rhythm analysis (NPCRA) indices included inter-daily stability (IS), intra-daily variability (IV), L5, M10, onset of L5 and M10, and relative amplitude.

### Human Intruder Test (HIT)

HIT was performed as previously described [60, 93]. Animals underwent a 10-min acclimation followed by a 2-min control recording. A non-threatening intruder then stood 0.3 m from the cage in profile without eye contact for 2 min, exited for 2 min, and returned for a 2-min threatening condition with direct eye contact. Sessions were video-recorded and analyzed by an observer blinded to group. Outcome measures included time spent in cage zones, including on the floor, low, middle or high zone of the cage, front, middle or back of the cage, vocalizations (tse-eggs), head and body bobs and locomotion.

### CatWalk XT gait analysis

Gait was quantified using CatWalk XT (Noldus Information Technology Inc.) on an illuminated glass walkway. Each animal performed 3 trials with 3 runs per trial (9 runs total). Qualified runs meeting quality criteria were selected for analysis. The primary outcome reported was percentage paw swing.

### Histology and immunohistochemistry

At 3 months post-transplantation, animals were euthanized by sodium pentobarbital overdose (100 mg/kg) followed by transcardiac perfusion with PBS followed by 4% paraformaldehyde. The brains were cryoprotected in an increasing gradient of sucrose solution (10, 20 and 30%), cryostat sectioned and processed for immunohistopathological analysis. Brain slices throughout the rostro-caudal extent were processed for hematoxylin and eosin (H&E) staining. Cell cultures were fixed with 4% paraformaldehyde for 15 min. Both cultured cells and brain sections were rinsed in PBS for 3x5 min then incubated for 2 hrs (cultures) or overnight (brain sections) with the appropriate primary antibodies for multiple labeling. Secondary antibodies raised in the appropriate hosts and conjugated to FITC, RITC, AMCA, CY3 or CY5 chromogenes (Jackson ImmunoResearch) were used. Cells and sections were counterstained with the nuclear marker 4’,6-diamidine-2’-phenylindole dihydrochloride (DAPI). Positive and negative controls were included in each run. Immunostained sections were coverslipped using fluorsave (Calbiochem) as the mounting medium. The following antibodies were used: Olig2 (monoclonal, 1:250, SantaCruz Biotechnology), β-tubulin Class-III (Monoclonal, 1:1000, Sigma-Aldrich, St. Louis, MO), Nestin (polyclonal, 1:100, EMD Millipore, Burlington, MA); SOX2 (polyclonal, 1:100, Abcam, Cambridge, United Kingdom); glial fibrillary acidic protein (GFAP, monoclonal, 1:1000, Chemicon; polyclonal 1:200, Aves Labs). Fluorescence was detected, analyzed, and photographed with a Zeiss LSM800 laser scanning confocal photomicroscope

## References

1. Rao V, Lyketsos C. Neuropsychiatric sequelae of traumatic brain injury. Psychosomatics. 2000;41(2):95–103. Epub 2000/04/06. doi: 10.1176/appi.psy.41.2.95. PubMed PMID: 10749946.

2. Moore EL, Terryberry-Spohr L, Hope DA. Mild traumatic brain injury and anxiety sequelae: a review of the literature. Brain Inj. 2006;20(2):117–32. Epub 2006/01/20. doi: 10.1080/02699050500443558. PubMed PMID: 16421060.

3. Yu S, Kaneko Y, Bae E, Stahl CE, Wang Y, van Loveren H, et al. Severity of controlled cortical impact traumatic brain injury in rats and mice dictates degree of behavioral deficits. Brain Res. 2009;1287:157–63. Epub 2009/07/04. doi: 10.1016/j.brainres.2009.06.067. PubMed PMID: 19573519.

4. Sanders MJ, Dietrich WD, Green EJ. Cognitive function following traumatic brain injury: effects of injury severity and recovery period in a parasagittal fluid-percussive injury model. J Neurotrauma. 1999;16(10):915–25. Epub 1999/11/05. PubMed PMID: 10547100.

5. Kimbler DE, Murphy M, Dhandapani KM. Concussion and the adolescent athlete. J Neurosci Nurs. 2011;43(6):286–90. Epub 2011/11/03. doi: 10.1097/JNN.0b013e31823858a6. PubMed PMID: 22045196.

6. The Management of Concussion/mTBI Working Group. VA/DoD Clinical Practice Guideline for Management of Concussion/Mild Traumatic Brain Injury. J Rehabil Res Dev. 2009;46(6):Cp1–68. Epub 2010/01/30. PubMed PMID: 20108447.

7. Faul M, Wald MM, Rutland-Brown W, Sullivent EE, Sattin RW. Using a cost-benefit analysis to estimate outcomes of a clinical treatment guideline: testing theBrain Trauma Foundation guidelines for the treatment of severe traumatic brain injury. J Trauma. 2007;63(6):1271–8. Epub 2008/01/24. doi: 10.1097/TA.0b013e3181493080. PubMed PMID: 18212649.

8. Menon DK, Schwab K, Wright DW, Maas AI. Position statement: definition of traumatic brain injury. Archives of physical medicine and rehabilitation. 2010;91(11):1637–40. Epub 2010/11/04. doi: 10.1016/j.apmr.2010.05.017. PubMed PMID: 21044706.

9. Carlson PJ, Singh JB, Zarate CA, Jr., Drevets WC, Manji HK. Neural circuitry and neuroplasticity in mood disorders: insights for novel therapeutic targets. Neurorx. 2006;3(1):22–41. Epub 2006/02/24. doi: 10.1016/j.nurx.2005.12.009. PubMed PMID: 16490411; PubMed Central PMCID: PMC3593361.

10. Adelson PD, Dixon CE, Robichaud P, Kochanek PM. Motor and cognitive functional deficits following diffuse traumatic brain injury in the immature rat. J Neurotrauma. 1997;14(2):99–108. Epub 1997/02/01. PubMed PMID: 9069441.

11. Brody DL, Mac Donald C, Kessens CC, Yuede C, Parsadanian M, Spinner M, et al. Electromagnetic controlled cortical impact device for precise, graded experimental traumatic brain injury. J Neurotrauma. 2007;24(4):657–73. Epub 2007/04/19. doi: 10.1089/neu.2006.0011. PubMed PMID: 17439349; PubMed Central PMCID: PMC2435168.

12. Bryant RA, O’Donnell ML, Creamer M, McFarlane AC, Clark CR, Silove D. The psychiatric sequelae of traumatic injury. Am J Psychiatry. 2010;167(3):312–20. Epub 2010/01/06. doi: 10.1176/appi.ajp.2009.09050617. PubMed PMID: 20048022.

13. Chauhan NB, Gatto R, Chauhan MB. Neuroanatomical correlation of behavioral deficits in the CCI model of TBI. J Neurosci Methods. 2010;190(1):1–9. Epub 2010/04/14. doi: 10.1016/j.jneumeth.2010.04.004. PubMed PMID: 20385166.

14. Jones NC, Cardamone L, Williams JP, Salzberg MR, Myers D, O’Brien TJ. Experimental traumatic brain injury induces a pervasive hyperanxious phenotype in rats. J Neurotrauma. 2008;25(11):1367–74. Epub 2008/12/09. doi: 10.1089/neu.2008.0641. PubMed PMID: 19061380.

15. Kennedy JE, Jaffee MS, Leskin GA, Stokes JW, Leal FO, Fitzpatrick PJ. Posttraumatic stress disorder and posttraumatic stress disorder-like symptoms and mild traumatic brain injury. J Rehabil Res Dev. 2007;44(7):895–920. Epub 2007/12/14. PubMed PMID: 18075948.

16. Rodgers KM, Bercum FM, McCallum DL, Rudy JW, Frey LC, Johnson KW, et al. Acute neuroimmune modulation attenuates the development of anxiety-like freezing behavior in an animal model of traumatic brain injury. J Neurotrauma. 2012;29(10):1886–97. Epub 2012/03/23. doi: 10.1089/neu.2011.2273. PubMed PMID: 22435644; PubMed Central PMCID: PMC3390983.

17. Schouten JW, Fulp CT, Royo NC, Saatman KE, Watson DJ, Snyder EY, et al. A review and rationale for the use of cellular transplantation as a therapeutic strategy for traumatic brain injury. Journal of Neurotrauma. 2004;21(11):1501–38. PubMed PMID: 15684646.

18. Schwarzbold ML, Rial D, De Bem T, Machado DG, Cunha MP, dos Santos AA, et al. Effects of traumatic brain injury of different severities on emotional, cognitive, and oxidative stress-related parameters in mice. J Neurotrauma. 2010;27(10):1883–93. Epub 2010/07/24. doi: 10.1089/neu.2010.1318. PubMed PMID: 20649482.

19. Washington PM, Forcelli PA, Wilkins T, Zapple DN, Parsadanian M, Burns MP. The effect of injury severity on behavior: a phenotypic study of cognitive and emotional deficits after mild, moderate, and severe controlled cortical impact injury in mice. J Neurotrauma. 2012;29(13):2283–96. Epub 2012/05/31. doi: 10.1089/neu.2012.2456. PubMed PMID: 22642287; PubMed Central PMCID: PMC3430487.

20. Pandey DK, Yadav SK, Mahesh R, Rajkumar R. Depression-like and anxiety-like behavioural aftermaths of impact accelerated traumatic brain injury in rats: a model of comorbid depression and anxiety? Behav Brain Res. 2009;205(2):436–42. Epub 2009/08/08. doi: 10.1016/j.bbr.2009.07.027. PubMed PMID: 19660499.

21. Koliatsos VE, Cernak I, Xu L, Song Y, Savonenko A, Crain BJ, et al. A mouse model of blast injury to brain: initial pathological, neuropathological, and behavioral characterization. J Neuropathol Exp Neurol. 2011;70(5):399–416. Epub 2011/04/14. doi: 10.1097/NEN.0b013e3182189f06. PubMed PMID: 21487304.

22. Ouellet MC, Beaulieu-Bonneau S, Morin CM. Sleep-wake disturbances after traumatic brain injury. Lancet Neurol. 2015;14(7):746–57. doi: 10.1016/s1474-4422(15)00068-x. PubMed PMID: 26067127.

23. Faul M, Coronado V. Chapter 1 - Epidemiology of traumatic brain injury. In: Grafman J, Salazar AM, editors. Handbook of Clinical Neurology. 127: Elsevier; 2015. p. 3–13.

24. Wilson L, Stewart W, Dams-O’Connor K, Diaz-Arrastia R, Horton L, Menon DK, et al. The chronic and evolving neurological consequences of traumatic brain injury. Lancet Neurol. 2017;16(10):813–25. Epub 20170912. doi: 10.1016/s1474-4422(17)30279-x. PubMed PMID: 28920887; PubMed Central PMCID: PMCPMC9336016.

25. Johnson VE, Stewart W, Smith DH. Axonal pathology in traumatic brain injury. Exp Neurol. 2013;246:35–43. Epub 2012/01/31. doi: 10.1016/j.expneurol.2012.01.013. PubMed PMID: 22285252.

26. Werner C, Engelhard K. Pathophysiology of traumatic brain injury. British Journal of Anaesthesia. 2007;99(1):4–9. doi: 10.1093/bja/aem131.

27. Chang F, Lee K, Ozturk ED, Chanfreau-Coffinier C, Merritt VC. Effects of TBI history and sleep on subjective cognition in Veterans: A VA Million Veteran Program study. J Psychiatr Res. 2025;193:112–9. Epub 20251114. doi: 10.1016/j.jpsychires.2025.11.005. PubMed PMID: 41273924.

28. McGee R, Montoya MA, Barber J, Joyner KJ, Nelson LD, Temkin N, et al. The Interaction of Sleep and Mood During Recovery from Mild Traumatic Brain Injury. Neurotrauma Rep. 2025;6(1):824–37. Epub 20250916. doi: 10.1177/2689288x251377033. PubMed PMID: 41220702; PubMed Central PMCID: PMCPMC12599806.

29. Cotter C, Tapp ZM, Ren C, Houle S, Mitsch J, Sheridan J, et al. Sleep fragmentation intensifies sleep architecture disruption and fatigue after traumatic brain injury. Exp Neurol. 2026;396:115544. Epub 20251105. doi: 10.1016/j.expneurol.2025.115544. PubMed PMID: 41202867.

30. Alvaro PK, Roberts RM, Harris JK. A Systematic Review Assessing Bidirectionality between Sleep Disturbances, Anxiety, and Depression. Sleep. 2013;36(7):1059–68. Epub 20130701. doi: 10.5665/sleep.2810. PubMed PMID: 23814343; PubMed Central PMCID: PMCPMC3669059.

31. Wickwire EM, Schnyer DM, Germain A, Williams SG, Lettieri CJ, McKeon AB, et al. Sleep, Sleep Disorders, and Circadian Health following Mild Traumatic Brain Injury in Adults: Review and Research Agenda. J Neurotrauma. 2018;35(22):2615–31. Epub 20180824. doi: 10.1089/neu.2017.5243. PubMed PMID: 29877132; PubMed Central PMCID: PMCPMC6239093.

32. Montgomery MC, Baylan S, Gardani M. Prevalence of insomnia and insomnia symptoms following mild-traumatic brain injury: A systematic review and meta-analysis. Sleep Med Rev. 2022;61:101563. Epub 20211102. doi: 10.1016/j.smrv.2021.101563. PubMed PMID: 35033968.

33. Cinotti R, Derouin Y, Chenet A, Oujamaa L, Glize B, Launey Y, et al. Standardized Outcomes for Randomized Controlled Trials Targeting Early Interventions in Patients With Moderate-to-Severe Traumatic Brain Injury: Protocol for the Development of a Core Outcome Set. JMIR Res Protoc. 2025;14:e54525. Epub 20250109. doi: 10.2196/54525. PubMed PMID: 39787594; PubMed Central PMCID: PMCPMC11757975.

34. Shaw JS, Woodard K, Krieg A, Bryant BR, Kentis S, Esagoff AI, et al. The impact of traumatic brain injury on sleep and associated neuroimaging changes: A systematic review. Sleep Med Rev. 2025;84:102155. Epub 20250911. doi: 10.1016/j.smrv.2025.102155. PubMed PMID: 41032947.

35. Roberts SSH, Owen PJ, Warmington SA, Trevenen J, Caeyenberghs K, McDonald SJ, et al. A systematic review and meta-analysis of sleep following mild traumatic brain injury: A synthesis of the literature according to age and time-since-injury. Sleep Med Rev. 2025;81:102072. Epub 20250218. doi: 10.1016/j.smrv.2025.102072. PubMed PMID: 40347689.

36. Ganesh A, Al-Shamli S, Mahadevan S, Chan MF, Burke DT, Al Rasadi K, et al. The Frequency of Neuropsychiatric Sequelae After Traumatic Brain Injury in the Global South: A systematic review and meta-analysis. Sultan Qaboos Univ Med J. 2024;24(2):161–76. Epub 20240527. doi: 10.18295/squmj.9.2023.056. PubMed PMID: 38828259; PubMed Central PMCID: PMCPMC11139369.

37. Marion D, Bullock MR. Current and future role of therapeutic hypothermia. J Neurotrauma. 2009;26(3):455–67. Epub 2009/03/19. doi: 10.1089/neu.2008.0582. PubMed PMID: 19292697.

38. Liao GP, Harting MT, Hetz RA, Walker PA, Shah SK, Corkins CJ, et al. Autologous bone marrow mononuclear cells reduce therapeutic intensity for severe traumatic brain injury in children. Pediatr Crit Care Med. 2015;16(3):245–55. doi: 10.1097/pcc.0000000000000324. PubMed PMID: 25581630; PubMed Central PMCID: PMCPMC4351120.

39. Cox CS. Safety of Autologous Stem Cell Treatment for Traumatic Brain Injury in Children (Clinical Trial Registration No. NCT00254722). clinicaltrials.gov. https://clinicaltrials.gov/study/NCT00254722. 2020.

40. Sharma A, Sane H, Kulkarni P, Yadav J, Gokulchandran N, Biju H, et al. Cell therapy attempted as a novel approach for chronic traumatic brain injury - a pilot study. Springerplus. 2015;4:26. Epub 20150117. doi: 10.1186/s40064-015-0794-0. PubMed PMID: 25628985; PubMed Central PMCID: PMCPMC4303601.

41. Cox CS, Jr., Baumgartner JE, Harting MT, Worth LL, Walker PA, Shah SK, et al. Autologous bone marrow mononuclear cell therapy for severe traumatic brain injury in children. Neurosurgery. 2011;68(3):588–600. doi: 10.1227/NEU.0b013e318207734c. PubMed PMID: 21192274.

42. Badner A, Cummings BJ. The endogenous progenitor response following traumatic brain injury: a target for cell therapy paradigms. Neural Regen Res. 2022;17(11):2351–4. doi: 10.4103/1673-5374.335833. PubMed PMID: 35535870; PubMed Central PMCID: PMCPMC9120693.

43. Sharp DJ, Scott G, Leech R. Network dysfunction after traumatic brain injury. Nat Rev Neurol. 2014;10(3):156–66. Epub 20140211. doi: 10.1038/nrneurol.2014.15. PubMed PMID: 24514870.

44. Armstrong RC, Sullivan GM, Perl DP, Rosarda JD, Radomski KL. White matter damage and degeneration in traumatic brain injury. Trends Neurosci. 2024;47(9):677–92. Epub 20240810. doi: 10.1016/j.tins.2024.07.003. PubMed PMID: 39127568.

45. Leskinen S, Mehta NH, Shah HA, Quelle M, Woodworth R, Dituri G, et al. Functional Connectivity Changes in Traumatic Brain Injury: A Systematic Review and Coordinate-Based Meta-Analysis of fMRI Studies. Neurology. 2025;105(9):e214298. Epub 20251017. doi: 10.1212/wnl.0000000000214298. PubMed PMID: 41105904.

46. Simon DW, McGeachy MJ, Bayır H, Clark RSB, Loane DJ, Kochanek PM. The far-reaching scope of neuroinflammation after traumatic brain injury. Nat Rev Neurol. 2017;13(9):572. Epub 20170804. doi: 10.1038/nrneurol.2017.116. PubMed PMID: 28776601.

47. Okonkwo DO, McAllister P, Achrol AS, Karasawa Y, Kawabori M, Cramer SC, et al. Mesenchymal Stromal Cell Implants for Chronic Motor Deficits After Traumatic Brain Injury: Post Hoc Analysis of a Randomized Trial. Neurology. 2024;103(7):e209797. Epub 20240904. doi: 10.1212/wnl.0000000000209797. PubMed PMID: 39231380; PubMed Central PMCID: PMCPMC11373674.

48. Badner A, Reinhardt EK, Nguyen TV, Midani N, Marshall AT, Lepe CA, et al. Freshly Thawed Cryobanked Human Neural Stem Cells Engraft within Endogenous Neurogenic Niches and Restore Cognitive Function after Chronic Traumatic Brain Injury. J Neurotrauma. 2021;38(19):2731–46. Epub 20210831. doi: 10.1089/neu.2021.0045. PubMed PMID: 34130484.

49. Beretta S, Cunningham KM, Haus DL, Gold EM, Perez H, Lopez-Velazquez L, et al. Effects of Human ES-Derived Neural Stem Cell Transplantation and Kindling in a Rat Model of Traumatic Brain Injury. Cell Transplant. 2017;26(7):1247–61. Epub 2017/09/22. doi: 10.1177/0963689717714107. PubMed PMID: 28933218; PubMed Central PMCID: PMCPMC5657732.

50. Haus DL, Lopez-Velazquez L, Gold EM, Cunningham KM, Perez H, Anderson AJ, et al. Transplantation of human neural stem cells restores cognition in an immunodeficient rodent model of traumatic brain injury. Exp Neurol. 2016;281:1–16. Epub 2016/04/16. doi: 10.1016/j.expneurol.2016.04.008. PubMed PMID: 27079998.

51. FDA US. Manufacturing Changes and Comparability for Human Cellular and Gene Therapy Products. Draft Guidance for Industry 2023:1–22.

52. FDA US. Potency Assurance for Cellular and Gene Therapy Products. Draft Guidance for Industry 2023:1–25.

53. FDA US. Considerations for the Use of Human- and Animal-Derived Materials in the Manufacture of Cell and Gene Therapy and Tissue-Engineered Medical Products Draft Guidance for Industry 2024:1–16.

54. Haus DL, Nguyen HX, Gold EM, Kamei N, Perez H, Moore HD, et al. CD133-enriched Xeno-Free human embryonic-derived neural stem cells expand rapidly in culture and do not form teratomas in immunodeficient mice. Stem Cell Res. 2014;13(2):214–26. Epub 20140710. doi: 10.1016/j.scr.2014.06.008. PubMed PMID: 25082219; PubMed Central PMCID: PMCPMC5675021.

55. Dixon CE, Clifton GL, Lighthall JW, Yaghmai AA, Hayes RL. A controlled cortical impact model of traumatic brain injury in the rat. J Neurosci Methods. 1991;39(3):253–62. Epub 1991/10/01. PubMed PMID: 1787745.

56. Lighthall JW. Controlled cortical impact: a new experimental brain injury model. J Neurotrauma. 1988;5(1):1–15. doi: 10.1089/neu.1988.5.1. PubMed PMID: 3193461.

57. Malloy KE, Li J, Choudhury GR, Torres A, Gupta S, Kantorak C, et al. Magnetic Resonance Imaging-Guided Delivery of Neural Stem Cells into the Basal Ganglia of Nonhuman Primates Reveals a Pulsatile Mode of Cell Dispersion. Stem Cells Transl Med. 2017;6(3):877–85. Epub 2017/03/16. doi: 10.5966/sctm.2016-0269. PubMed PMID: 28297573; PubMed Central PMCID: PMCPMC5442780.

58. McEntire CR, Choudhury GR, Torres A, Steinberg GK, Redmond DE, Jr., Daadi MM. Impaired Arm Function and Finger Dexterity in a Nonhuman Primate Model of Stroke: Motor and Cognitive Assessments. Stroke. 2016;47(4):1109–16. Epub 2016/03/10. doi: 10.1161/strokeaha.115.012506. PubMed PMID: 26956259.

59. Choudhury GR, Daadi MM. Charting the onset of Parkinson-like motor and non-motor symptoms in nonhuman primate model of Parkinson’s disease. PLoS One. 2018;13(8):e0202770. Epub 2018/08/24. doi: 10.1371/journal.pone.0202770. PubMed PMID: 30227600; PubMed Central PMCID: PMCPMC6107255 alter our adherence to PLOS ONE policies on sharing data and materials.

60. Santangelo AM, Ito M, Shiba Y, Clarke HF, Schut EH, Cockcroft G, et al. Novel Primate Model of Serotonin Transporter Genetic Polymorphisms Associated with Gene Expression, Anxiety and Sensitivity to Antidepressants. Neuropsychopharmacology. 2016;41(9):2366–76. Epub 20160321. doi: 10.1038/npp.2016.41. PubMed PMID: 26997299; PubMed Central PMCID: PMCPMC4946067.

61. Mallya S, Sutherland J, Pongracic S, Mainland B, Ornstein TJ. The manifestation of anxiety disorders after traumatic brain injury: a review. J Neurotrauma. 2015;32(7):411–21. Epub 20150123. doi: 10.1089/neu.2014.3504. PubMed PMID: 25227240.

62. McKee AC, Cantu RC, Nowinski CJ, Hedley-Whyte ET, Gavett BE, Budson AE, et al. Chronic traumatic encephalopathy in athletes: progressive tauopathy after repetitive head injury. J Neuropathol Exp Neurol. 2009;68(7):709–35. Epub 2009/06/19. doi: 10.1097/NEN.0b013e3181a9d503. PubMed PMID: 19535999; PubMed Central PMCID: PMC2945234.

63. McKee AC, Stein TD, Nowinski CJ, Stern RA, Daneshvar DH, Alvarez VE, et al. The spectrum of disease in chronic traumatic encephalopathy. Brain. 2013;136(Pt 1):43–64. Epub 2012/12/05. doi: 10.1093/brain/aws307. PubMed PMID: 23208308; PubMed Central PMCID: PMC3624697.

64. O’Donnell ML, Creamer M, Pattison P, Atkin C. Psychiatric morbidity following injury. Am J Psychiatry. 2004;161(3):507–14. Epub 2004/03/03. PubMed PMID: 14992977.

65. Zatzick DF, Jurkovich GJ, Gentilello L, Wisner D, Rivara FP. Posttraumatic stress, problem drinking, and functional outcomes after injury. Arch Surg. 2002;137(2):200–5. Epub 2002/03/05. PubMed PMID: 11822960.

66. Gavett BE, Stern RA, McKee AC. Chronic traumatic encephalopathy: a potential late effect of sport-related concussive and subconcussive head trauma. Clin Sports Med. 2011;30(1):179–88, xi. Epub 2010/11/16. doi: 10.1016/j.csm.2010.09.007. PubMed PMID: 21074091; PubMed Central PMCID: PMC2995699.

67. Bendlin BB, Ries ML, Lazar M, Alexander AL, Dempsey RJ, Rowley HA, et al. Longitudinal changes in patients with traumatic brain injury assessed with diffusion-tensor and volumetric imaging. Neuroimage. 2008;42(2):503–14. Epub 20080507. doi: 10.1016/j.neuroimage.2008.04.254. PubMed PMID: 18556217; PubMed Central PMCID: PMCPMC2613482.

68. Sidaros A, Skimminge A, Liptrot MG, Sidaros K, Engberg AW, Herning M, et al. Long-term global and regional brain volume changes following severe traumatic brain injury: a longitudinal study with clinical correlates. Neuroimage. 2009;44(1):1–8. Epub 20080904. doi: 10.1016/j.neuroimage.2008.08.030. PubMed PMID: 18804539.

69. Wu TC, Wilde EA, Bigler ED, Li X, Merkley TL, Yallampalli R, et al. Longitudinal changes in the corpus callosum following pediatric traumatic brain injury. Dev Neurosci. 2010;32(5-6):361–73. Epub 20101014. doi: 10.1159/000317058. PubMed PMID: 20948181; PubMed Central PMCID: PMCPMC3073757.

70. Brezova V, Moen KG, Skandsen T, Vik A, Brewer JB, Salvesen O, et al. Prospective longitudinal MRI study of brain volumes and diffusion changes during the first year after moderate to severe traumatic brain injury. NeuroImage Clinical. 2014;5:128–40. Epub 20140328. doi: 10.1016/j.nicl.2014.03.012. PubMed PMID: 25068105; PubMed Central PMCID: PMCPMC4110353.

71. Cox CS, Jr., Notrica DM, Juranek J, Miller JH, Triolo F, Kosmach S, et al. Autologous bone marrow mononuclear cells to treat severe traumatic brain injury in children. Brain. 2024;147(5):1914–25. doi: 10.1093/brain/awae005. PubMed PMID: 38181433; PubMed Central PMCID: PMCPMC11068104.

72. Spitz G, Maller JJ, O’Sullivan R, Ponsford JL. White matter integrity following traumatic brain injury: the association with severity of injury and cognitive functioning. Brain Topogr. 2013;26(4):648–60. Epub 20130327. doi: 10.1007/s10548-013-0283-0. PubMed PMID: 23532465.

73. Kinnunen KM, Greenwood R, Powell JH, Leech R, Hawkins PC, Bonnelle V, et al. White matter damage and cognitive impairment after traumatic brain injury. Brain. 2011;134(Pt 2):449–63. Epub 20101229. doi: 10.1093/brain/awq347. PubMed PMID: 21193486; PubMed Central PMCID: PMCPMC3030764.

74. Hulkower MB, Poliak DB, Rosenbaum SB, Zimmerman ME, Lipton ML. A decade of DTI in traumatic brain injury: 10 years and 100 articles later. AJNR American journal of neuroradiology. 2013;34(11):2064–74. Epub 20130110. doi: 10.3174/ajnr.A3395. PubMed PMID: 23306011; PubMed Central PMCID: PMCPMC7964847.

75. Farbota KD, Bendlin BB, Alexander AL, Rowley HA, Dempsey RJ, Johnson SC. Longitudinal diffusion tensor imaging and neuropsychological correlates in traumatic brain injury patients. Front Hum Neurosci. 2012;6:160. Epub 20120619. doi: 10.3389/fnhum.2012.00160. PubMed PMID: 22723773; PubMed Central PMCID: PMCPMC3378081.

76. Hutchinson EB, Schwerin SC, Avram AV, Juliano SL, Pierpaoli C. Diffusion MRI and the detection of alterations following traumatic brain injury. J Neurosci Res. 2018;96(4):612–25. Epub 20170613. doi: 10.1002/jnr.24065. PubMed PMID: 28609579; PubMed Central PMCID: PMCPMC5729069.

77. Choi JY, Hart T, Whyte J, Rabinowitz AR, Oh SH, Lee J, et al. Myelin water imaging of moderate to severe diffuse traumatic brain injury. NeuroImage Clinical. 2019;22:101785. Epub 20190316. doi: 10.1016/j.nicl.2019.101785. PubMed PMID: 30927603; PubMed Central PMCID: PMCPMC6444291.

78. Churchill NW, Caverzasi E, Graham SJ, Hutchison MG, Schweizer TA. White matter during concussion recovery: Comparing diffusion tensor imaging (DTI) and neurite orientation dispersion and density imaging (NODDI). Human brain mapping. 2019;40(6):1908–18. Epub 20181226. doi: 10.1002/hbm.24500. PubMed PMID: 30585674; PubMed Central PMCID: PMCPMC6865569.

79. Wright AD, Jarrett M, Vavasour I, Shahinfard E, Kolind S, van Donkelaar P, et al. Myelin Water Fraction Is Transiently Reduced after a Single Mild Traumatic Brain Injury- -A Prospective Cohort Study in Collegiate Hockey Players. PLoS One. 2016;11(2):e0150215. Epub 20160225. doi: 10.1371/journal.pone.0150215. PubMed PMID: 26913900; PubMed Central PMCID: PMCPMC4767387.

80. Hillary FG, Grafman JH. Injured Brains and Adaptive Networks: The Benefits and Costs of Hyperconnectivity. Trends Cogn Sci. 2017;21(5):385–401. Epub 20170401. doi: 10.1016/j.tics.2017.03.003. PubMed PMID: 28372878; PubMed Central PMCID: PMCPMC6664441.

81. Caeyenberghs K, Leemans A, Leunissen I, Gooijers J, Michiels K, Sunaert S, et al. Altered structural networks and executive deficits in traumatic brain injury patients. Brain Struct Funct. 2014;219(1):193–209. Epub 20121212. doi: 10.1007/s00429-012-0494-2. PubMed PMID: 23232826.

82. Todd BP, Chimenti MS, Luo Z, Ferguson PJ, Bassuk AG, Newell EA. Traumatic brain injury results in unique microglial and astrocyte transcriptomes enriched for type I interferon response. J Neuroinflammation. 2021;18(1):151. Epub 20210705. doi: 10.1186/s12974-021-02197-w. PubMed PMID: 34225752; PubMed Central PMCID: PMCPMC8259035.

83. Arneson D, Zhang G, Ahn IS, Ying Z, Diamante G, Cely I, et al. Systems spatiotemporal dynamics of traumatic brain injury at single-cell resolution reveals humanin as a therapeutic target. Cell Mol Life Sci. 2022;79(9):480. Epub 20220811. doi: 10.1007/s00018-022-04495-9. PubMed PMID: 35951114; PubMed Central PMCID: PMCPMC9372016.

84. Swaro A, Bristow BN, Anwer M, Zhang AA, Kraus L, Gandhi RK, et al. Widespread and cell-type-specific transcriptomic reorganization following mild traumatic brain injury. Cell Rep. 2025;44(6):115795. Epub 20250605. doi: 10.1016/j.celrep.2025.115795. PubMed PMID: 40478731.

85. Koupourtidou C, Schwarz V, Aliee H, Frerich S, Fischer-Sternjak J, Bocchi R, et al. Shared inflammatory glial cell signature after stab wound injury, revealed by spatial, temporal, and cell-type-specific profiling of the murine cerebral cortex. Nat Commun. 2024;15(1):2866. Epub 20240403. doi: 10.1038/s41467-024-46625-w. PubMed PMID: 38570482; PubMed Central PMCID: PMCPMC10991294.

86. Dockree PM, Robertson IH. Electrophysiological markers of cognitive deficits in traumatic brain injury: a review. Int J Psychophysiol. 2011;82(1):53–60. Epub 20110114. doi: 10.1016/j.ijpsycho.2011.01.004. PubMed PMID: 21238506.

87. Morelli N, Johnson NF, Kaiser K, Andreatta RD, Heebner NR, Hoch MC. Resting state functional connectivity responses post-mild traumatic brain injury: a systematic review. Brain Inj. 2021;35(11):1326–37. Epub 20210906. doi: 10.1080/02699052.2021.1972339. PubMed PMID: 34487458.

88. NINDS/NIH. Rethinking TBI Classification for Clinical Care and Research https://wwwnindsnihgov/sites/default/files/documents/Rethinking%20TBI%20Classification%20for%20Clinical%20Care%20and%20Research_26June%202023pdf? 2024:1–27.

89. Ng K, Mikulis DJ, Glazer J, Kabani N, Till C, Greenberg G, et al. Magnetic resonance imaging evidence of progression of subacute brain atrophy in moderate to severe traumatic brain injury. Archives of physical medicine and rehabilitation. 2008;89(12 Suppl):S35–44. doi: 10.1016/j.apmr.2008.07.006. PubMed PMID: 19081440.

90. NIH/NICHD. NIH Consensus Development Conference on Rehabilitation of Persons with Traumatic Brain Injury https://wwwnichdnihgov/publications/pubs/TBI_1999/NIH_Consensus_Statement. 1999:1–21.

91. Daadi MM, Hu S, Klausner J, Li Z, Sofilos M, Sun G, et al. Imaging neural stem cell graft-induced structural repair in stroke. Cell Transplant. 2013;22(5):881–92. Epub 2012/10/10. doi: 10.3727/096368912x656144. PubMed PMID: 23044338.

92. Daadi MM, Li Z, Arac A, Grueter BA, Sofilos M, Malenka RC, et al. Molecular and magnetic resonance imaging of human embryonic stem cell-derived neural stem cell grafts in ischemic rat brain. Mol Ther. 2009;17(7):1282–91. PubMed PMID: 19436269.

93. Agustín-Pavón C, Braesicke K, Shiba Y, Santangelo AM, Mikheenko Y, Cockroft G, et al. Lesions of ventrolateral prefrontal or anterior orbitofrontal cortex in primates heighten negative emotion. Biol Psychiatry. 2012;72(4):266–72. Epub 20120412. doi: 10.1016/j.biopsych.2012.03.007. PubMed PMID: 22502990.

